# Structural Basis for Near Infrared Absorption and Noncanonical Photoisomerization in Neorhodopsin

**DOI:** 10.64898/2026.10.08.757671

**Authors:** Kirill Kovalev, Ben Boehlke, Probal Nag, Maximilian Große, Anastasia Kraskov, Sagie Katz, Peter Hildebrandt, Franz Bartl, Igor Schapiro, Peter Hegemann, Matthias Broser

## Abstract

In retinal proteins, light absorption leads to a double bond isomerization of the chromophore, followed by conformational relaxations that drive functional protein rearrangements. The balance between photoisomerization and radiative or non-radiative decays of the excited state is adjusted to the demands of the respective photoreceptor function. The molecular determinants controlling the excited state processes remain poorly understood and are the focus of this combined crystallographic, spectroscopic, and theoretical work on the recently discovered neorhodopsin (NeoR). NeoR exhibits a highly red-shifted and fluorescent dark state and selectively photoisomerizes from all-*trans* to 7-*cis*. We show that the local electrostatic environment reduces the negative charge density near the Schiff base and suppresses torsional motion around the canonical C13=C14 isomerization site of microbial rhodopsins. Instead, the chromophore-binding pocket permits only low-efficiency C7=C8 isomerization, accompanied by polyene inversion, β-ionone ring displacement, and Schiff base deprotonation. Consequently, unlike other retinal photoreceptors, NeoR undergoes structural changes localized near the β-ionone ring.

## Introduction

In photoreceptors such as rhodopsins or phytochromes light absorption by the chromophore leads to a double bond isomerization as the starting point of the photoinduced reaction cascades that eventually lead to a biological response^1,2^. In these chromophores, electronic excitation triggers charge redistribution perturbing the interactions with the immediate protein environment. These interactions shape the energetic landscape for bond twisting and determine the efficiencies of the various channels for the excited state decay, i.e. the non-productive fluorescence and internal conversion, and the productive photoisomerization. The balance between steric constraints imposed by the protein scaffold and electrostatic interactions arising from the local polarity governs both the quantum yield and directionality of the isomerization process. Understanding this interplay is the basis for rationalizing how protein environments tune chromophore function.

Retinal proteins are particularly appropriate photoreceptors to address this issue since there is already a large body of experimental and theoretical data on the photochemical and photophysical processes of the chromophore, a protonated retinylidene Schiff base (RSBH^+^) ^1,3–8^. Furthermore, the recent discovery of neorhodopsins (NeoRs), a rhodopsin module in fungal heterodimeric rhodopsin–guanylyl cyclases (Supplementary Fig. 1), has challenged the paradigm that rhodopsins are weakly fluorescent and photoisomerize at the C11=C12 (animal rhodopsins) or C13=C14 double bond (microbial rhodopsins) ^9^. NeoR absorbs in the near-infrared (NIR) region around 700 nm (Fig. 1A), far beyond the most red-shifted animal and microbial rhodopsins ^10,11^. This extreme opsin shift is accompanied by an unusually long excited-state lifetime and strong fluorescence with a quantum yield of approximately 20%, which is at least three orders of magnitude higher than those typically observed in other rhodopsins (0.001– 0.01%) ^12^. Unlike canonical rhodopsins, continuous illumination of NeoR promotes only low-efficiency photoisomerization into an unusual 7-*cis* isomer with deprotonated Schiff base and an absorption maximum in the near UV ^13,14^. Notably, NeoR is the only known retinal protein that selectively stabilizes this sterically constrained 7-*cis* configuration, that otherwise only occurs as a marginal byproduct. It is widely accepted that the photochemical and photophysical properties of the protonated retinylidene Schiff base (RSB) chromophore are governed by steric constraints and electrostatic interactions. Thus, elucidation of their specific contributions in NeoR is essential for understanding the molecular basis of its unusual excited state behavior in contrast to canonical retinal proteins.

**Fig. 1.**
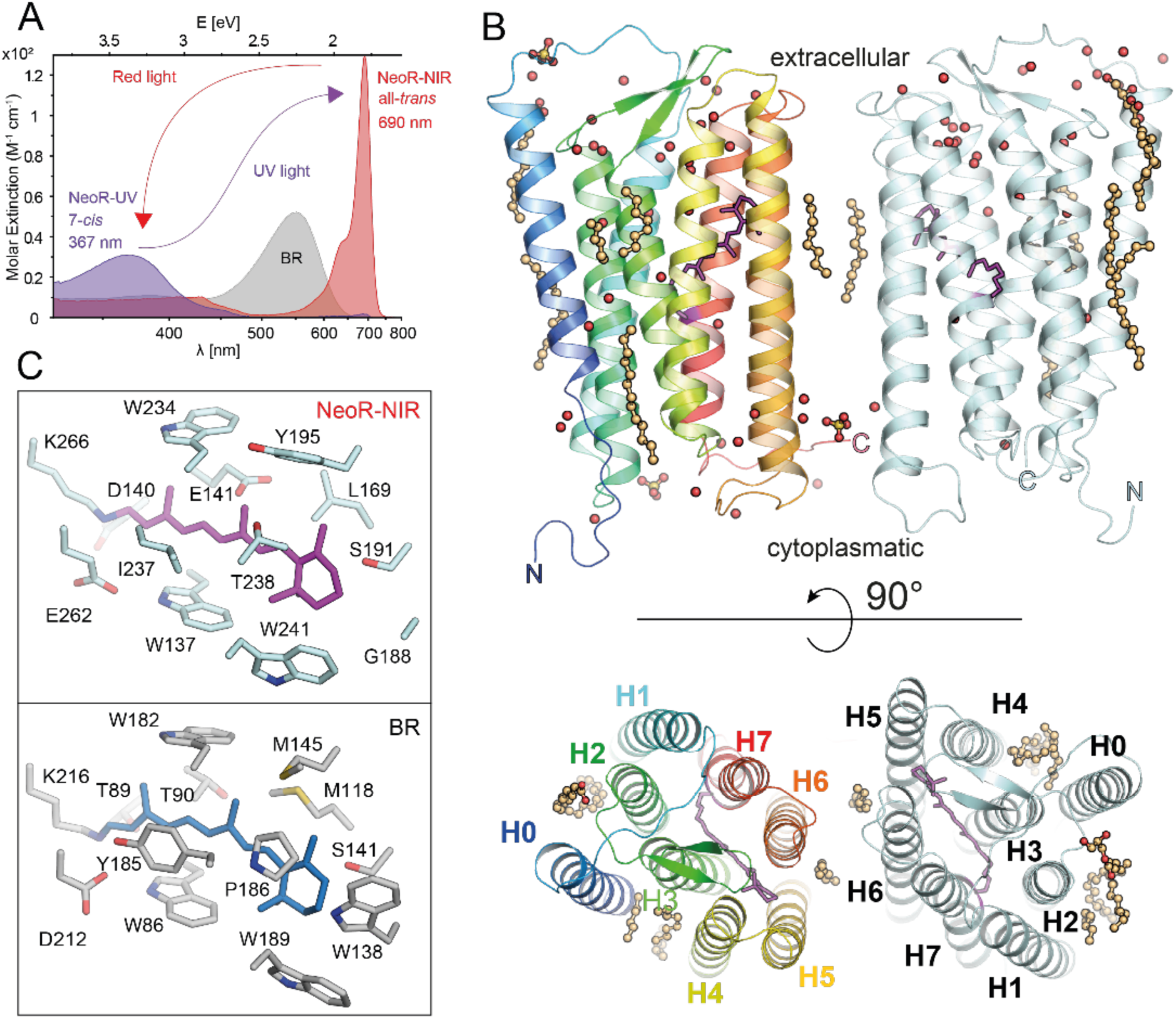
Spectral properties and structural overview of RgNeoR. (A) Molar extinction spectra of *R. globosum* neorhodopsin: NeoR-NIR (red)/NeoR-UV (violet) state, and prototypical microbial rhodopsin (bacteriorhodopsin (BR (gray))) molar extinction is plotted versus a linear energy scale (eV) corresponding to a log wavelength scale (nm). (B) Overall structure of the NeoR-NIR rhodopsin homodimer (aa 1-279), with water (red spheres) lipids (orange ball-and-sticks) and retinylidene (magenta). (C) The chromophore binding pocket of NeoR-NIR (upper panel) and BR (lower panel, pdb: 1QHJ ^42^) with nearby side chains drawn as sticks.

So far structural insight into NeoR has relied upon sequence analysis and homology modeling ^9,15^. In this work, however, we succeeded in determining the crystal structure of the NeoR-module in the NIR-absorbing dark and UV-absorbing UV state, harboring the all-*trans* and 7-*cis* chromophore, respectively. Both states were characterized by spectroscopic techniques and the dark state structure served as a basis for multiscale simulations of absorption and photoisomerization.

## Results

### Crystal structures of NeoR-NIR and NeoR-UV

We crystallized the detergent-purified rhodopsin domain of NeoR (residues 1-279) from *R. globosum* (“NeoR” in the following) in lipidic cubic phase (LCP). Crystals formed at pH 6.0 and displayed a vivid pale green color indicative for the NeoR-NIR state and diffracted up to a resolution of 2.1 Å (Supplementary Fig. 2 A, B; Supplementary Table 1). In addition, we determined the 2.4 Å structure of NeoR-NIR at room temperature using serial crystallography (SX; Supplementary Table 1), which supported that the structural features associated with near-infrared light absorption are present at both cryo and ambient temperatures. The asymmetric unit contains a homodimer, consistent with a previous low-resolution single-particle cryo-EM structure ^16^, but with an helix H6-rather than helix H5-mediated interface, suggesting structural rearrangements during crystallization. Neither arrangement is physiologically relevant, as NeoR functions exclusively within a heterodimeric rhodopsin–guanylyl cyclase complex (Supplementary Fig. 1).

Except for the N-terminal residues 1-15 in monomer A and 1-17 in monomer B, all residues of the protein are well-resolved, allowing for accurate modeling of side chain conformations. Additionally, several lipids, three sulfate ions, and 68 protein bound water molecules are clearly defined. Most water molecules are found in an extracellular cavity defined by helices H3, 4, and 5 (Fig. 1 B). Furthermore, we identified two strongly bound Zn²⁺ ions in each of the two NeoR monomers, originating from yeast as the host organism (Supplementary Fig. 3).

We illuminated the crystals with far-red light (660 nm) over several minutes and observed complete loss of color, consistent with a quantitative conversion to NeoR-UV (Supplementary Fig. 2 E, F). Single crystals of the photoconverted protein showed limited diffraction, and an additional SX approach was applied to improve the NeoR-UV structural dataset to 2.5 Å resolution (Supplementary Table 1). Importantly, the above-mentioned Zn2+ ions bound to NeoR also mediate crystal contacts; therefore, to further improve diffraction data quality we soaked protein crystals in precipitant solution supplemented with 10 mM zinc acetate. Then, using SX, we obtained an additional structure of NeoR-UV at 2.4 Å (Supplementary Table 1). Both reported NeoR-UV structures show similar rearrangements of the chromophore and its binding pocket upon photoconversion.

### The structure of the NeoR-NIR state

The overall architecture of NeoR resembles that of canonical microbial rhodopsins, comprising seven transmembrane helices that surround the retinylidene chromophore. As in other enzyme-rhodopsins, this core is supplemented by an additional N-terminal helix (H0; Fig. 1 B). Compared with rhodopsin-phosphodiesterase (Rh-PDE;^17^), helix H0 in NeoR is slightly shifted toward helix H3. The protonated Schiff base is surrounded by three carboxylates (E136, D140, and E262), referred to as the active site EDE triad. A ∼25 Å–long continuous hydrogen-bond chain exists parallel to the membrane plane, involving residues S166, H167, N32, N143, H139, and three water molecules, ultimately connecting the EDE triad with E141 (Fig. 2 A). These lateral polar interactions are only found in NeoR and may contribute to reduced conformational dynamics around the chromophore by fixing the relative orientation of the helices involved. On the proximal side, the EDE triad is bridged by a water molecule with R133 (Fig. 2 B), which further provides a tight structural connection between adjacent helices H1, H3, and H7.

**Fig. 2.**
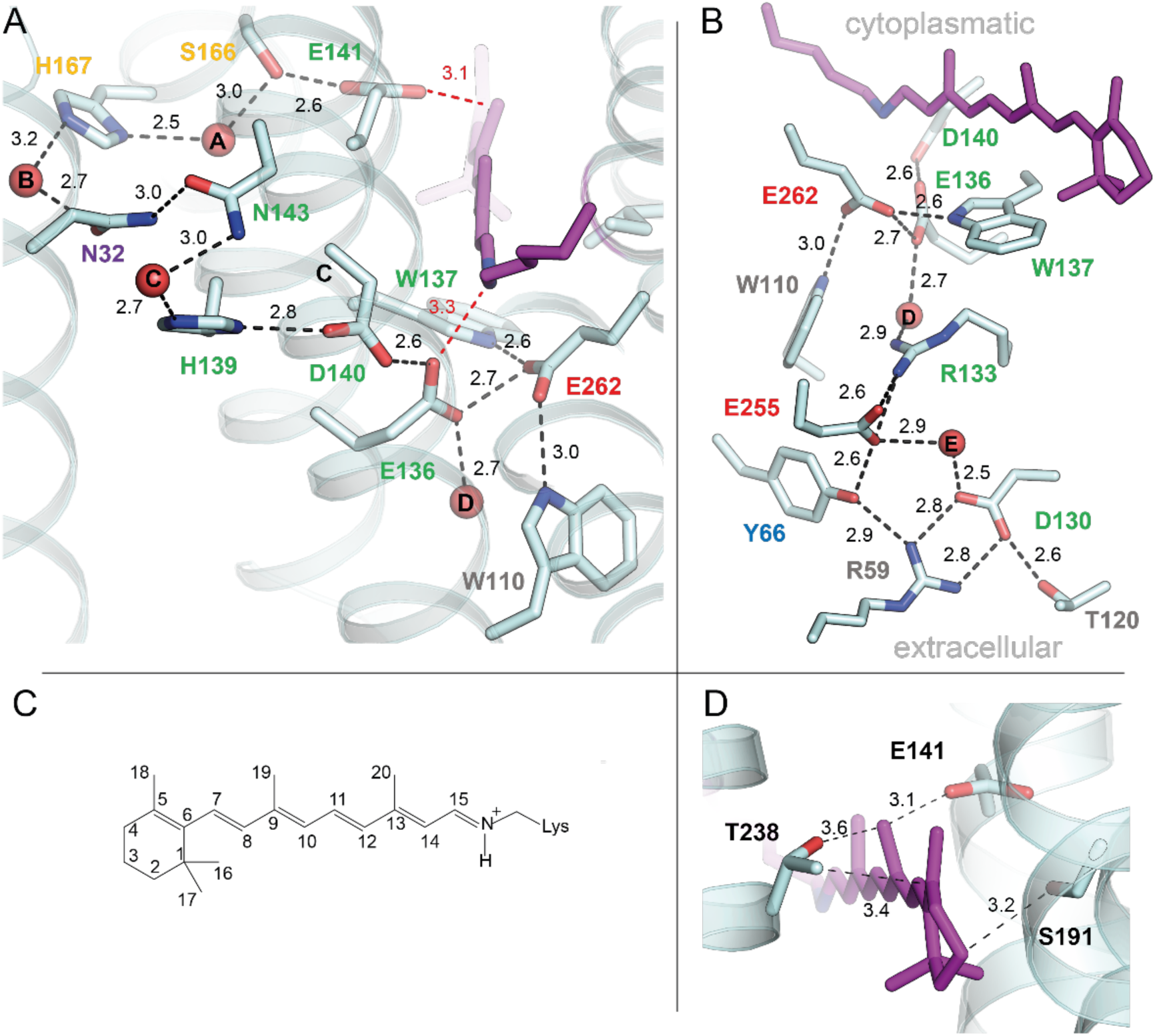
Hydrogen-bond networks connected to the chromophore binding pocket. Expanded horizontal (A) and vertical (B) hydrogen-bond network in NeoR-NIR. Residues are colored according to their helices: H0 (violet), H1 (blue), H2 (gray), H3 (green), H4 (orange), H7 (red). Red dashed line indicates closest distance of carboxylates to the retinal (distances in Å). (C) *all-trans* retinylidene structure and atom numbering. (D) The all*-trans* retinylidene chromophore in NeoR-NIR showing bent C18 and C19 methyl groups.

Continuous electron density for the NeoR-NIR retinylidene enabled unambiguous determination of the chromophore in the all-*trans* configuration (Fig. 3 A). The retinal binding pocket is evolutionary inherited from ancestral fungal rhodopsin guanylyl-cyclases ^18^ with many amino acids conserved in the majority of microbial rhodopsins like the two tryptophans W137 and W234 that keep the retinylidene in place (Fig. 1 C). In contrast, the conserved Tyr/Phe position adjacent to the polyene chain is substituted by Ile (I237), creating additional space between the chromophore and its surrounding protein matrix (Fig. 1 C and S4). The β-ionone ring is in a 6-s-*trans* conformation and in close contact with S191 and T238 (Fig. 2 D), two residues that facilitate red-shifted absorption in canonical microbial rhodopsins ^19^. The retinylidene C18 and C19 methyl groups are twisted out of the chromophore’s conjugation plane (Fig. 2 D) due to steric strains by the proximity of T238 and E141, respectively. The latter residue was considered as a key element for the NIR absorption in a computational study based on a homology modelling ^15^.

**Fig. 3.**
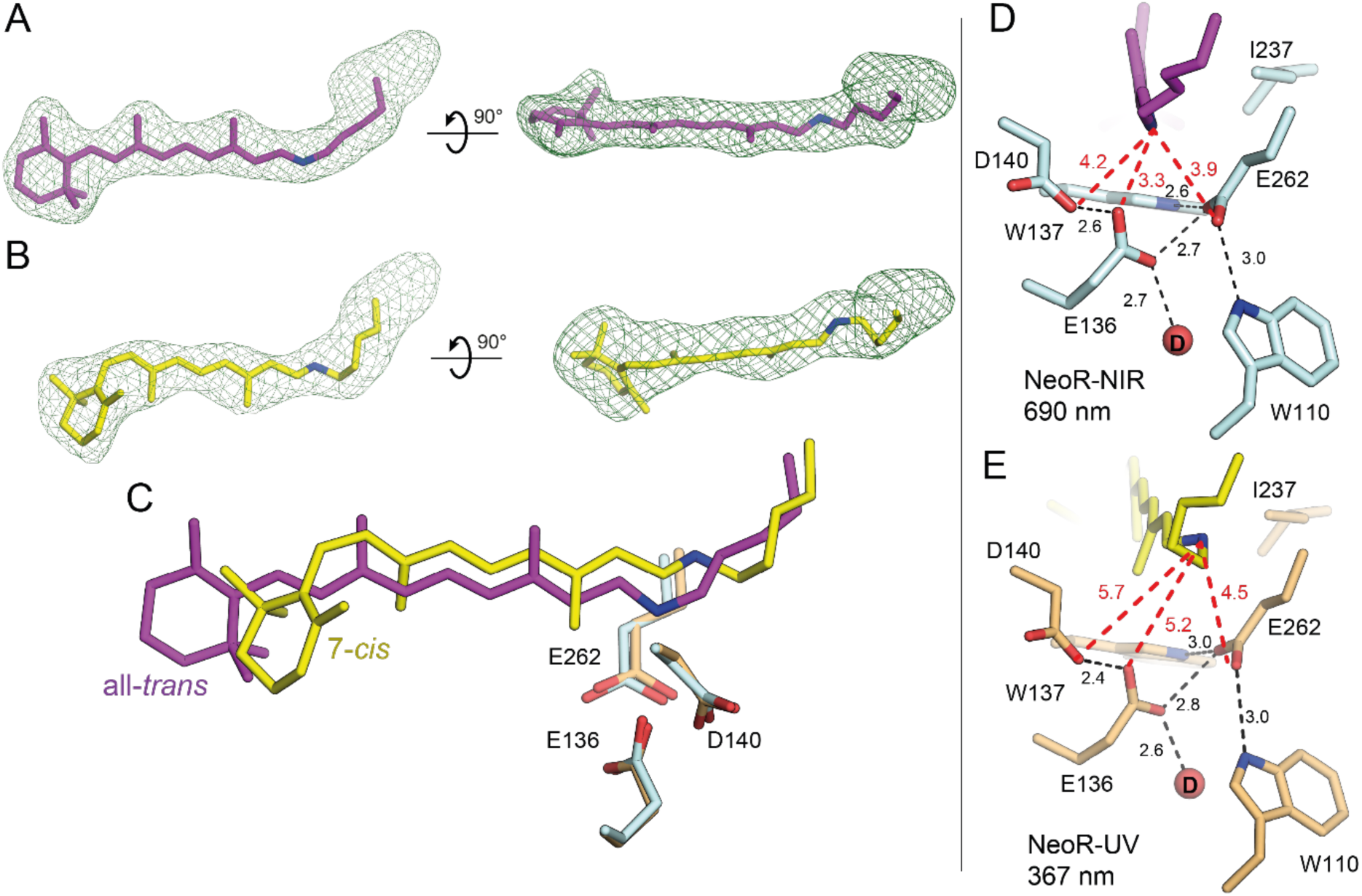
The chromophore structure in NeoR-NIR and NeoR-UV. (A,B) Polder maps of the retinylidene cofactor in the NIR-absorbing (A) and UV-absorbing (B) state of NeoR contoured at the level of 3.0σ. (C) Superposition of the chromophore in NeoR-UV (lime; sidechains: orange) and NeoR-NIR (magenta; sidechains: cyan). Note the inversion and displacement of the RSB nitrogen but minor displacement of the active site residues. (D,E) Close-up of the RSB active site in NeoR-NIR (D) and UV state (E), red-dashed lines indicate the closest distance of carboxylic acids side chains of the EDE-triad (E136, D140, E262) to the RSB nitrogen. All distances in Å.

The immediate Schiff base environment of NeoR-NIR (Fig. 3 D) has been considered as the main determinant for the far-red absorption ^9^. The EDE triad forms a tightly packed carboxylate network but exhibits only weak, if any, coupling to the protonated RSBH⁺ nitrogen, with O:::N distances of 3.3, 4.2, and 3.9 Å for E136, D140, and E262, respectively (Fig. 3 D). These distances exceed those typically observed in other microbial rhodopsins (Supplementary Fig. 5), and no bridging water molecules are detected. The RSBH⁺ nitrogen is oriented towards E262, whose side chain is further stabilized in its position by the tryptophans W110 (3.0 Å) and W137 (2.6 Å). The lack of stabilizing interactions promotes the delocalization of the positive Schiff base charge over the entire chromophore.

The protonation state of the EDE-triad carboxylic acids has been already previously discussed ^9,20^. From mutagenesis studies we have concluded that D140 is protonated, whereas several quantum mechanics/molecular mechanics (QM/MM) calculations based on homology models suggested that, in addition to D140, either E136 or E262, or both are also protonated ^9,15^ The experimental structure in this work shows the arrangement of the EDE-triad, E141, and other residues that differ from previous models. Therefore, we recalculated the excitation energy for all possible protonation patterns of the EDE triad. Only models with protonated D140 and E262 showed minimal deviations in the EDE triad arrangement relative to the crystal structure and close agreement with the NeoR-NIR absorption spectrum whereas protonation of E136 is still left ambiguous. For both protonated and deprotonated E136, the predicted excitation energies and simulated spectra closely match the experimental data including a pronounced shoulder (Supplementary Table 2 and Supplementary Fig. 6 A, respectively) that reflects strong coupling between vibrational and electronic transition. The similar-spectra of the protonated and deprotonated E136 systems arises from comparable charge distributions in the protonated Schiff base due to almost identical electrostatic environments experienced by the chromophore (Supplementary Fig. 6 B).

### The structure of the NeoR-UV state

For NeoR-UV, the maximum of the absorption spectrum and the resonance Raman band at 1566 cm^−1^ and the lack of H/D isotopic shifts (Supplementary Fig. 7) indicate a deprotonated retinylidene chromophore ^21,22^. Retinal extraction experiments on NeoR from *R. globosum* provided evidence for a 7-*cis* isomer, consistent with the finding from fungus *O. mucronatum*.^13,14^ Indeed our structural data show that the C7=C8 bond adopts a *cis* configuration (Fig. 3 B,C). However, what was not anticipated is that the main part of the polyene chain between C8 and Cδ of Lys266 is flipped by ∼180°. Consequently, the polyene chain including the Schiff base nitrogen moves towards the extracellular surface by ∼1Å (Fig. 3 C). In addition, the structure shows an isomerization from 6-s-*trans* to 6-s-*cis* minimizing steric repulsion of the C16/C17 methyl groups with C19 methyl of the rotated polyene chain.

Despite the ∼1 Å movement of the RSB nitrogen upon formation of NeoR-UV, the location and conformation of the crucial EDE triad, and the tryptophans W110 and W137 appear unaffected (Figs. 3 C-E).

The FoUV-FoNIR isomorphous difference electron density map highlighted further structural rearrangements upon photoconversion (Fig. 4). The changes were primarily localized close to the β-ionone ring where isomerization of the C7=C8 bond is accompanied by a pronounced displacement of the β-ionone ring and nearby residues of H4 and H5, including E141 of H3 (Fig. 4 B). This rearrangement reduces the accessibility of the retinal binding pocket from the lipid bilayer via locking the fenestration between helices H5 and H6 (Fig. 4 D and Supplementary Fig. 8). Although a similar fenestration accommodates carotenoid antenna molecules in numerous proton-pumping rhodopsins ^23,24^, its role in NeoR remains unclear.

**Fig. 4.**
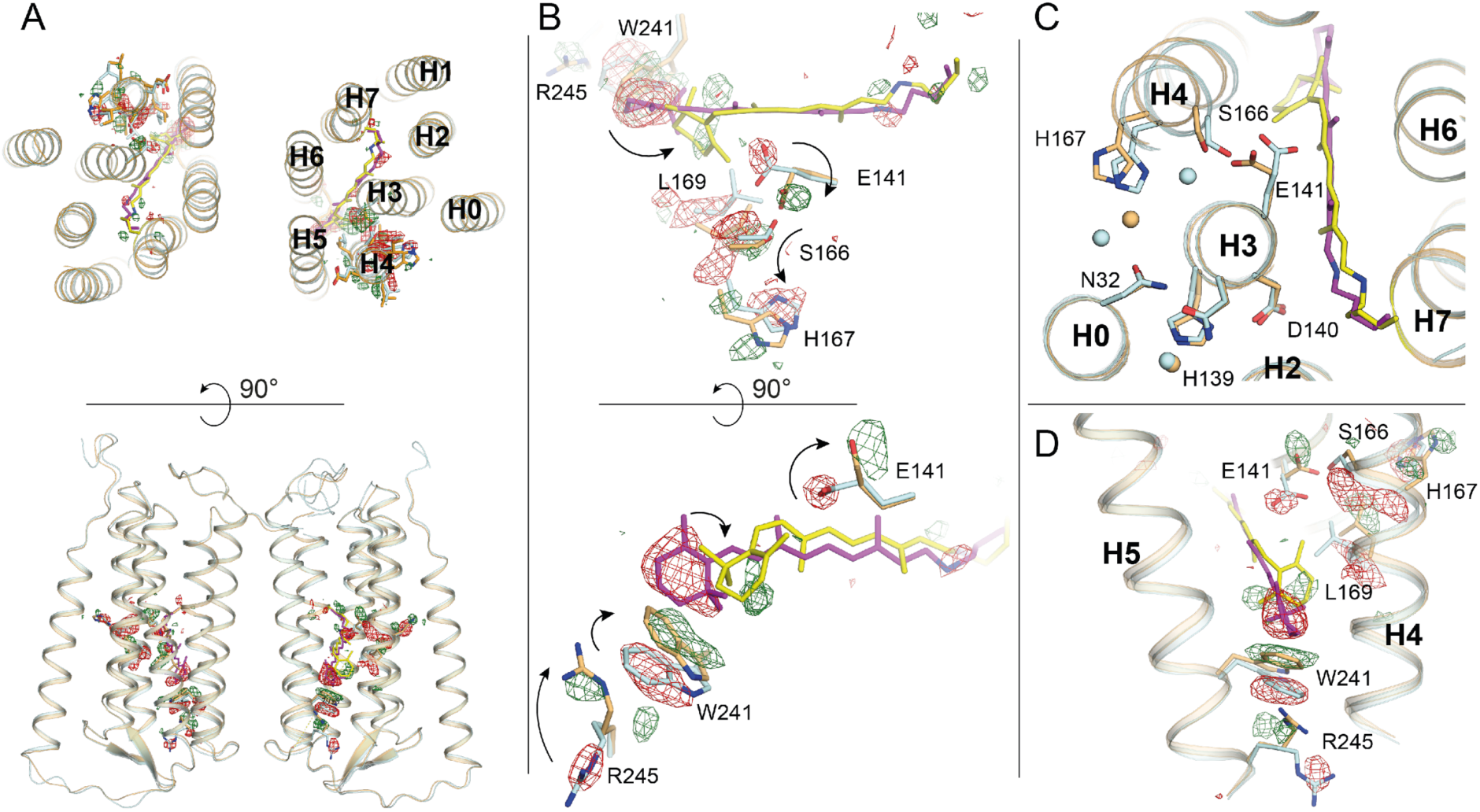
Structural changes induced by photoconversion. (A) Top and side view of the two NeoR protomers. FoUV-FoNIR map is contoured at the level of 3.0σ to visualize the changes localized near the β-ionone ring of the retinal and H4. (B) Detailed view of the retinylidene with NeoR-UV (lime, orange) overlayed with NeoR-NIR (magenta, cyan). Black arrows indicate the most pronounced rearrangements in NeoR upon the formation of the UV-absorbing state. (C) Top view on the hydrogen bond network around H3, showing the flipping of E141 towards S166 in NeoR-UV. (D) Close-up on conformational changes at the interface of H4 and H5.

In all previously studied microbial and animal rhodopsins, photoisomerization induces structural changes primarily near the RSB active site, often involving a change in the protonation state of the counterion complex due to repositioning of the RSB nitrogen from a polar into a non-polar environment after isomerization. The β-ionone ring, by contrast, typically undergoes only minimal spatial shifts. Thus, the all-*trans* to 7-*cis* photoisomerization in NeoR enables the transmission of conformational changes to a different region of the protein, far from the RSB.

Photoconversion to NeoR-UV triggers a rotation of E141 away from the polyene chain (Fig. 4 B,C), strengthening its interaction with S166-OH and reinforcing the connection between helices H3 and H4. While such coupling between these helices is known to accelerate thermal re-isomerization in many microbial rhodopsins ^25–27^ the noncanonical 7-*cis* configuration in NeoR-UV remains thermally stable. The conformational rearrangement of E141 is consistent with the NeoR-UV minus NeoR-NIR FTIR difference spectrum, which exhibits a negative signal at 1756 cm⁻¹ in the dark state that shifts to 1737 cm^−1^ upon photoproduct formation, indicating light-induced changes in the hydrogen-bond environment of protonated E141 (Supplementary Fig. 9). In D2O this band pattern is down-shifted to 1746 (-)/ 1728 (+) and is absent in the E141A mutant allowing a clear assignment to E141. Subsequently, the hydrogen bond network involving S166 and H167 changes, and the imidazole ring appears to undergo a 180° rotation, as inferred from shifts in nearby water positions (Fig. 4 B,C). Since the RSB is deprotonated in NeoR-UV, photoconversion must have triggered the transfer of the RSBH^+^ proton to another residue. The invariance of the EDE triad between NeoR-NIR and NeoR-UV argues against proton transfer to EDE as proximal acceptor (Fig. 3 E). Instead, the proton is likely relayed to more distal residues along the hydrogen-bonded network (Fig. 2 A).

### Blocked photoisomerization pathway in NeoR-NIR

NeoR differs from typical microbial rhodopsins in that it does not undergo photoisomerization to the 13-*cis* isomer. In contrast, NeoR exhibits strong fluorescence, typically attributed to a barrier preventing access to the conical intersection leading to prolonged excited-state lifetime ^9^. Based on the NeoR-NIR crystal structure, we performed hybrid quantum mechanics/molecular mechanics (QM/MM) simulations, to elucidate the origin of this unusual photochemistry. Because of the special role of the counterion complex we have included the EDE triad and E141 in the QM region, with only E136 deprotonated (see Methods section). An excited state (S1) relaxed potential energy scan was performed along the torsional coordinate of the C13=C14 bond (Fig. 5 B) and compared with bacteriorhodopsin (BR), the best studied microbial rhodopsin (Fig. 5 C). A relaxed scan was performed because nonadiabatic molecular dynamics simulations would be prohibitive to study the excited state of NeoR which is reported to be 1 ns. Both negatively charged counterions (D85 and D212) and one nearby water molecule (Wat401) were included in the QM partition of BR to account for the polarization effect around the Schiff base. In NeoR, the S1 energy increases monotonically with increasing twist of the C13=C14 bond (Fig. 5 B). In BR, by contrast, the energy decreases after an out-of-plane twist of 30°, reaching a conical intersection (CI) that mediates 13-*cis* formation. The S₁ energy profile of BR is strongly asymmetric, indicating significant pre-twist of the C13=C14 bond at the Franck-Condon (FC)/ground state geometry and a preferred clockwise rotation consistent with experiments ^28^. We note that although nonadiabatic dynamics would yield a more complete description of the excited-state relaxation, the present relaxed scans provide a comparative view of the accessibility of the C13=C14 torsional pathway in NeoR and BR.

**Fig. 5.**
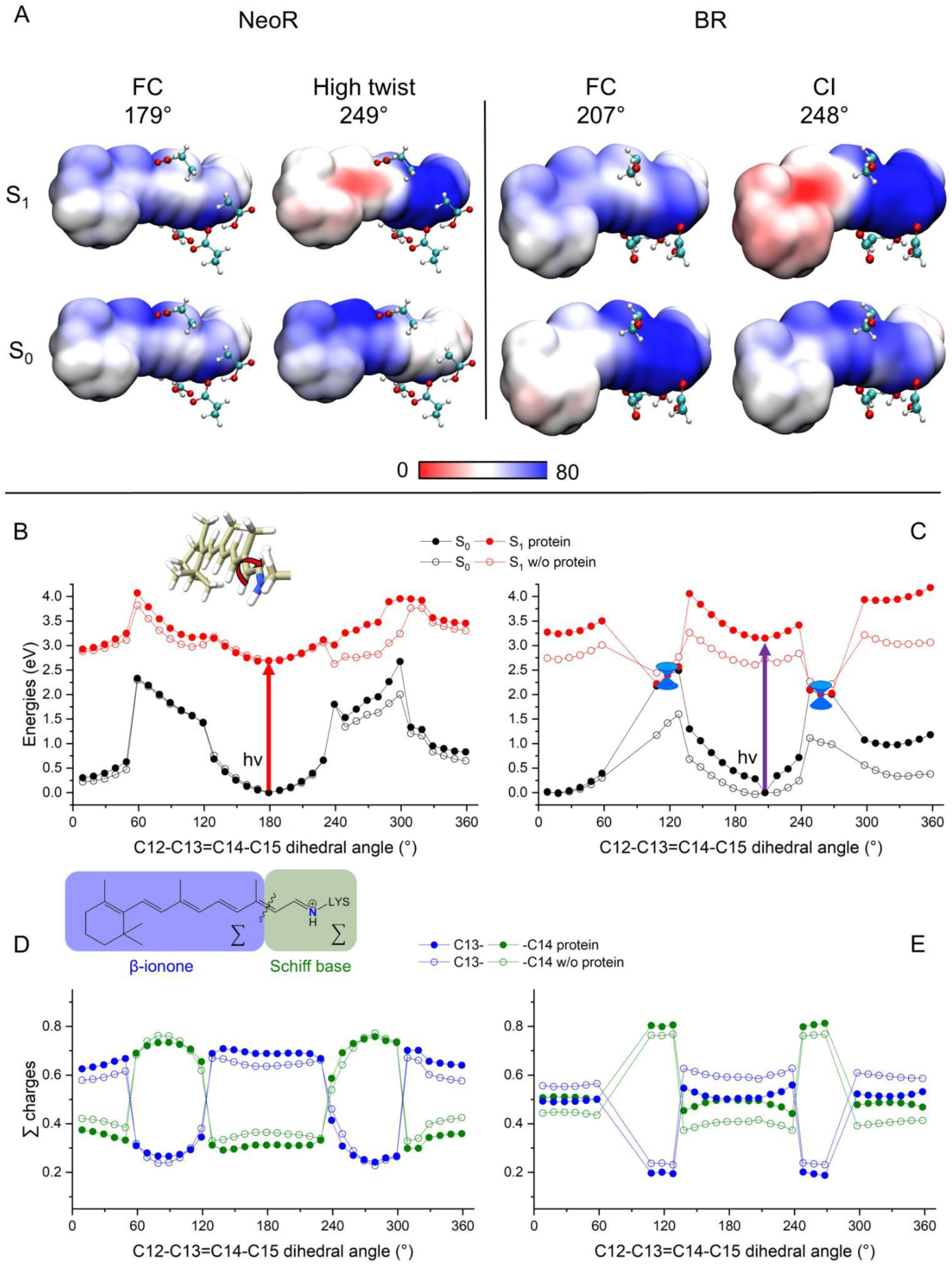
QM/MM simulations on NeoR-NIR and bacteriorhodopsin (BR). (A) Electrostatic potential (ESP) maps of retinal chromophore in the ground (S0) and excited (S1) states in a model with one negative counterion of NeoR (E136 deprotonated), and two negative charges in the case of BR. The ESP maps are computed for the Franck–Condon points and at geometries with high twist of the C13=C14 double bond. Selected residues are shown: NeoR: EDE-triad (E136, D140, E262) and E141. Br: D85, T90, D212 and water. (B, C) Relaxed potential energy scan along the C13=C14 double bond torsional coordinate in NeoR and BR, respectively. (D, E) S1 positive charge distribution (Mulliken analysis) along the Schiff base and β-ionone ring fragments of the chromophore during the relaxed scan in the two proteins, respectively. The bond length alternation (BLA) of the retinal chromophore calculated for each geometry along the relaxed scan is shown in orange. The energies and charge distributions are shown for the chromophore with and without the effects of the protein environment.

To sort out the electrostatic contribution of the protein environment, we calculated the energy profiles without the protein moiety, while keeping the chromophore geometry fixed. Under these conditions, the energy evolution in BR becomes similar to that in NeoR, and no conical intersection is observed in either protein (Fig. 5 B,C). This highlights the essential role of protein electrostatics, particularly the counterions, in enabling the non-radiative decay pathway.

To rationalize these differences, we analyzed the charge distribution along the retinal chromophore. In BR, the positive charge is mainly near the Schiff base in the ground state but shifts toward the β-ionone ring upon excitation (Fig. 5 A). NeoR displays a markedly different pattern: the positive charge is more delocalized along the polyene chain, with substantial density near the C9 methyl group and the β-ionone ring in both the ground and excited states. In BR, the negatively charged residues D85 and D212 stabilize the positive charge in the ground state. In NeoR, E136 forms strong interactions with the neighboring residues D140 and E262 within the EDE triad. This local network reduces the effective electrostatic interaction between E136 and the protonated Schiff base, resulting in a weaker counterion–Schiff base interaction than observed in canonical microbial rhodopsins such as BR.

To examine the influence of the protein environment on isomerization, we monitored the S1 charge distribution during the rotation around the C13=C14 bond. For this analysis the sum of partial charges in S₁ (Mulliken used in (Fig. 5) and Hirshfeld used in (Supplementary Fig. 10)) was calculated for two fragments ^29^: a β-ionone fragment extending to C13 (named C13-fragment in the following) and a Schiff base C14-fragment ranging from C14 to the iminium group (Fig. 5 D,E). In NeoR, the positive charge in S1 remains largely localized on the C13-fragment (Fig. 5 D). In BR, the charge is more evenly distributed between the two fragments in S1 (Charge transfer (CT) state, Fig. 5 E and Supplementary Fig. 10). When protein electrostatics are removed, the calculated charge distribution for BR becomes NeoR-like, confirming that the observed differences arise from the electrostatic environment and not from retinal geometry.

Twisting the C13=C14 double bond in NeoR does not change the charge distribution up to ∼50°. In contrast, in BR the positive charge progressively shifts toward the β-ionone ring enhancing the CT-character ^30^. At larger distortions, however, in both systems disruption of π-conjugation leads to a twist-induced intramolecular charge transfer (TICT) back to the C14-fragment, a charge distribution similar to the S0 state of BR (Fig. 5 A, D, E). This change in charge distribution is not accompanied by an abrupt change in the retinal bond length pattern. But, despite these similarities, a conical intersection is reached only in case of BR. This clear distinction can be traced back to differences in the counterion environment near the protonated Schiff base. In BR the negatively charged D85 and D212 effectively stabilize the positive charge on the Schiff base fragment, lowering the S1 energy, whereas NeoR lacks comparable stabilization.

Finally, we performed a relaxed scan around the C7=C8 bond in NeoR. In this case the chromophore approaches to a conical intersection region at a dihedral twist of approximately 60° (Supplementary Fig. 12). Because this bond lies at the distal end of the chromophore and the positive charge is distributed along a polyene chain, only a small fraction of the charge moves during the rotation towards the Schiff base even at highly twisted geometries (Supplementary Fig. 12). The electrostatic influence of the counterion complex is therefore much weaker compared to other double bond isomerizations. However, the photoisomerization of the C7=C8 bond requires a rotation of the entire polyene chain (Fig. 3), which is unfavorable and explains its low yield. A complete dynamical description of this low-yield pathway remains challenging due to the coupled retinal rearrangements accompanying C7=C8 isomerization. Both protonated and deprotonated E136 models predict similar energy profiles for C13=C14 and C7=C8 double bond torsions, reflecting the underlying similarity in the electrostatics (Supplementary Fig. 6).

## Discussion

Our combined crystallographic, spectroscopic, and multiscale quantum chemical approach reveals how the protein environment of NeoR fundamentally reshapes the photochemistry of the protonated retinal Schiff base chromophore. The crystal structures of the NeoR-NIR dark state and the NeoR-UV photoproduct show that light-induced structural changes are localized primarily near the β-ionone ring rather than around the Schiff base region, in striking contrast to canonical microbial and animal rhodopsins.

The NeoR-UV photoproduct reveals that, in addition to the unusual C7=C8 isomerization reported recently ^13^, the displacement of the β-ionone ring is accompanied by isomerization of the C6-C7 single bond and a flip of the polyene chain.

Hybrid QM/MM simulations based on the experimentally determined NeoR-NIR structure provide a mechanistic explanation for this unusual photochemistry. Protonation and interactions within the EDE triad weaken its effective electrostatic interaction with the positively charged chromophore. As a result, the positive charge is less localized at the Schiff base nitrogen and becomes more delocalized along the retinal polyene chain. This reduced electrostatic stabilization of the ground state lowers the energy gap between the ground and excited states compared with other microbial rhodopsins, thereby enabling NIR absorption.

Recent computational studies came to similar conclusions, although they relied on homology models of NeoR ^9,14,15^. Despite structural differences, the overall electronic structure of the chromophore appears to be consistently reproduced across models featuring reduced negative countercharge near the Schiff base, in agreement with the experimentally observed NeoR Raman spectra. This electrostatic environment also affects the access to the conical intersection associated with the canonical C13=C14 isomerization.

In the excited state of BR, the rotation of the C13=C14 bond causes the positive charge to shift further toward the β-ionone ring. This observation is in line with BR and bovine rhodopsin literature ^30–33^. At large distortions, however, our simulations reveal an abrupt change in charge distribution towards the Schiff base region near the conical intersection that has not been proposed previously. A related twist-induced charge localization near the Schiff base has recently been reported for engineered Arch rhodopsins, where it modulates the barrier to photoisomerization ^34–36^. In bacteriorhodopsin, the positive charge can be stabilized by the two negatively charged counterions, which favors completion of the torsional motion. In NeoR by contrast, this stabilization is absent, making rotation around the C13=C14 bond energetically unfavorable.

The abrupt redistribution of charge close to the conical intersection appears to be essential for rationalizing the calculated energy profiles. Only such a rapid change in the electronic structure can account for the stabilization of the twisted geometry that enables access to the conical intersection in bacteriorhodopsin. The present calculations therefore describe the relative accessibility of the excited-state pathway rather than providing a rigorous characterization of conical intersection geometries. The TDA-CAM-B3LYP scans are therefore interpreted as comparative excited-state energy profiles rather than a direct description of conical intersection topology. The presence of this abrupt charge redistribution feature may also explain why our simulations differ from several previous theoretical studies, which reported a more gradual charge migration along the chromophore. In our calculations, the inclusion of the EDE triad and E141 in NeoR, as well as the two negatively charged counterions (D85 and D212) and a nearby water molecule in BR, within the QM region allows the electrostatic response of the active site to be captured more accurately, revealing a sharper redistribution of charge at large torsional distortions. In molecular dynamics simulations, such a region of the potential energy surface may only be visited for a very short time. Experimentally, such a transient charge rearrangement would be extremely difficult to detect, because it occurs only in the immediate vicinity of the conical intersection and therefore on an ultrafast timescale. As a result, the process may be too short-lived to be resolved by current spectroscopic techniques, which typically probe earlier stages of the excited-state dynamics ^37^.

To further test the role of counterions, we calculated the partial charges with fully protonated active-site carboxylates or replaced them by alanine in both systems (Supplementary Fig. 11). In NeoR, this results in the positive charge being largely localized on the β-ionone fragment at the Franck–Condon point. In both cases, the BR excited-state profile becomes similar to NeoR, with suppression of the low-energy C13=C14 torsional pathway, underscoring the critical importance of negative charge near the protonated Schiff base in modulating the photochemistry. This is in line with spectroscopic studies of microbial rhodopsins and hybrid QM/MM simulations of channelrhodopsin ^38–41^. Simulations of the NeoR E141A mutant yielded results similar to those of NeoR WT (Supplementary Fig. 11 C), consistent with the minor effect of this mutation on the photochemistry.

NeoR provides a natural example of how protein electrostatics can shift retinal photophysics from an ultrafast, nonradiative photoisomerization regime toward an almost non-reactive, highly fluorescent state. However, the chromophore retains access to a conical intersection along the more distant C7=C8 bond, where the influence of the counterion complex is significantly weaker. This still enables the low-efficiency noncanonical photoisomerization pathway observed experimentally. In principle, such a reaction coordinate could exist in other rhodopsins but is masked by the dominant C11=C12 or C13=C14 isomerization reactions in previously studied systems.

## Methods

### Protein expression and purification

To express the rhodopsin domain of NeoR from *Rhizoclosmatium globosum* (UniProtKB: A0A1Y2CSJ0; residues 1–279) in *Pichia pastoris*, a human codon–optimized NeoR DNA sequence was cloned into a pPICZ vector (Thermo Fisher Scientific, Waltham, MA, USA). The construct carried a C-terminal TEV protease recognition site followed by a StrepII tag. Transformation, selection of recombinant clones, and protein expression were carried out primarily according to the manufacturer’s instructions (Thermo Fisher Scientific), with minor modifications. Briefly, protein expression in buffered growth medium supplemented with 5 µM all-trans retinal was induced by the addition of 2.5% (v/v) methanol. Cells were harvested 24 h post-induction and disrupted using a high-pressure homogenizer (HTU Digi-French-Press, G. Heinemann, Germany). Cell debris was separated by centrifugation (16000 xg, 10 min, 4°C) and subsequently crude membranes were pelleted by ultracentrifugation (180000 xg, 45 min, 4°C). The recombinant rhodopsin was solubilized in 50 mM HEPES (pH 7.4), 100 mM NaCl, 1.5% (w/v) dodecylmaltoside (DDM; Glycon, Luckenwalde, Germany), and 0.3% (w/v) cholesteryl hemisuccinate (CHS; Sigma-Aldrich, Merck KGaA, Darmstadt, Germany) and purified by affinity chromatography on a 5 ml Strep-Tactin®XT Superflow® column (IBA GmbH, Göttingen, Germany). Colored fractions were pooled and further purified by size exclusion chromatography (16/600 Hiload with Superdex 200pg (Cytiva, Marlborough, MA, USA)). Final fractions for crystallization were concentrated in 25 mM HEPES (pH7.4), 50 mM NaCl, 0.02% DDM, 0.004% CHS using Amicon Ultra centrifugal filters (MWCO 100 kDa; Merck Millipore, Burlington, MA, USA). Protein production of the E141A mutant was carried out analogously to the wild type.

### Protein crystallization

The crystals of *Rg*NeoR were grown with an *in meso* approach. In particular, the solubilized protein (20 mg/ml) in the crystallization buffer was mixed with a premelted at 42°C monopalmitolein (Nu-Chek Prep) in a 2:3 ratio (protein:lipid) to form a lipidic mesophase. The mesophase was homogenized in coupled syringes (Hamilton) by transferring the mesophase from one syringe to another until a homogeneous and gel-like material was formed. Then, 100 nl drops of a protein–mesophase mixture were spotted on a 96-well LCP glass sandwich plate (Marienfeld) and overlaid with 400 nL of precipitant solution by means of the Mosquito crystallization robot (SPT Labtech). The initial crystals were obtained using in the D2 condition (0.1M Lithium sulfate, 0.1M MES pH 6.0, 30% PEG 400) of MEMMESO screen (Molecular Dimensions). This condition was optimized manually by screening various pH values and PEG 400 and lithium sulfate concentrations. The best crystals were obtained using 0.15 M Lithium sulfate, 0.1M MES pH 6.0, 40% PEG 400 as precipitant. The crystals were grown at 22°C and reached their final size of ∼20×10x5 µm^3^ in 2 weeks.

For the determination of the near-infrared-absorbing *Rg*NeoR structure at 100K, once the crystals reached their final size, crystallization wells were opened, and drops containing the protein-mesophase mixture were covered with 100 μl of the respective precipitant solution. For the data collection, harvested crystals were incubated for 5 min in the precipitant solution. Crystals were harvested using micromounts (Mitegen, USA), flash-cooled, and stored in liquid nitrogen. For the determination of the UV-absorbing *Rg*NeoR structure, crystallization wells were opened, and drops containing the protein-mesophase mixture were covered with 100 μl of the respective precipitant solution. In case of Zn^2+^-bound structure, the precipitant solution was supplemented with 10 mM zinc acetate. Then, the drop has been illuminated by 660 nm LED light for 10 minutes. The crystals were inspected using a microscope (Leica) to validate the photoconversion of the protein to the UV-absorbing state by the loss of characteristic cyan color. Crystals were then harvested using micromounts (Mitegen, USA), flash-cooled, and stored in liquid nitrogen. For the determination of the Zn^2+^-bound RgNeoR structure in the UV-absorbing state, crystals were harvested using micromeshes (Mitegen, USA) for serial crystallography data collection.

For the determination of the near-infrared-absorbing *Rg*NeoR structure at 293K, the crystals were grown in CrystalDirectTM (Mitegen, USA) plates using 0.1M MES pH 6.0, 40% PEG 400 as precipitant. 40 µl of the precipitant solution was dispensed in the mother liquor reservoirs of each well, 100 nl of the mesophase was used per drop covered by 400 nl of the precipitant solution.

### Diffraction and X-ray fluorescence data collection and treatment

X-ray diffraction data were collected at the P14 beamline of PETRAIII (Hamburg, Germany) using an EIGER2 X 16M CdTe detector. X-ray fluorescence spectra were acquired at beamline P13 using Amptek XR-100SDD energy-dispersive detector and PX5 digital pulse processor (Amptek, Inc., Bedford, MA, USA), using incident photon energy 12.7 keV. The data collection was performed using MxCube2 software. For the collection of the serial crystallography data using CrystalDirectTM plates, the plate was scanned across the focused X-ray beam (5×7 µm^2^, 1.3×10^12^ photons/s) with an oscillation of 0.001° per image using plate manipulator goniometer head (Arinax, France). Diffraction images from the traditional rotational data collection were processed using XDS. The reflection intensities were scaled and merged using the Staraniso server. Diffraction images from the serial crystallography data were processed using CrystFEL (v0.10.1) using the following parameters for indexamajig: --multi --peaks=peakfinder8 --threshold=5 --min-snr=5 --indexing=xgandalf --integration=rings-grad --int-radius=2,4,6 --highres=2.0. The reflection intensities were scaled and merged using partialator (model unity and number of iterations 1 with the final high resolution cutoff of 2.4 Å for the *Rg*NeoR structure in the near-infrared-absorbing state at 293K and 2.3 Å for the *Rg*NeoR structure in the UV-absorbing state at 100K). There is no possibility of twinning for the crystals. The data collection and treatment statistics are presented in Table S1.

### Crystal structure determination and refinement

Initial phases were successfully obtained in the P212121 space groups by molecular replacement using MOLREP from the CCP4 program suite using the model of monomeric *Rg*NeoR predicted by AlphaFold2. The initial model was iteratively refined using REFMAC5 and Coot. The structure refinement statistics are presented in Table S1. The difference Folight-Fodark and polder omit maps were built using the PHENIX program suite.

### IR-spectroscopy

NeoR WT and E141A samples were mounted by stepwise drying the protein solution on a BaF2 window under a stream of dry air, followed by subsequent rehydration. To achieve a homogeneous sample thickness, the sample was sealed by another BaF2 window with a 6 µm thick PTFE spacer placed between them. The measurements were performed after an equilibration time of at least 120 minutes at Room temperature. The sample was illuminated with a 685 nm or 400 nm laser diode for a duration of 240 seconds. All spectra were recorded at 2 cm^−1^ resolution using a Vertex 80v FTIR Spectrometer (Bruker Optics) equipped with a liquid-nitrogen-cooled mercury-cadmium-telluride (MCT) detector (Kolmar Technologies). The difference spectra were calculated and processed with the OPUS 7.5 software (Bruker Optics) by using the single-channel spectra before and after illumination of the sample.

### Resonance Raman spectroscopy

Resonance Raman spectra of NeoR-UV in H₂O and D₂O buffers were recorded with 413 nm excitation from a Kr⁺ ion laser (Coherent, Santa Clara, CA, USA). Raman signals were collected in a 180° backscattering geometry using a confocal LabRam HR spectrometer (Horiba, Villeneuve, France) equipped with a liquid-nitrogen-cooled CCD detector. The dispersion was 0.4 cm⁻¹ per pixel, corresponding to a spectral resolution of 1.2 cm⁻¹. Typical measurements used an accumulation time of 1 h and a laser power of ∼1 mW at the sample. Low-temperature experiments were performed with a Linkam cryostat (Linkam Scientific Instruments, Surrey, UK) mounted on a computer-controlled XY stage (OWIS GmbH, Germany).

### Computational Methodology

Hybrid quantum mechanics/molecular mechanics (QM/MM) simulations were performed using the crystal structure as the starting point. Due to similarity in the models, monomer A was used for the simulations. The amino acid residues were protonated according to pH 7 except for the four titratable residues around the retinal chromophore -E136, D140, E141 and E262. For these four residues, various protonation states were modelled, where the proton was added to either of the carboxylate oxygen atoms in cis/trans conformation. This was performed to understand the effect of these residues on the spectral tuning mechanism. The resulting models after protonation were optimized through a QM/MM approach using ChemShell interfaced with ORCA ^43,44^. The QM part includes the retinal and the lysine sidechain forming the protonated Schiff base along with the four titratable residues mentioned above. The QM/MM boundary was placed between Cα and Cβ atoms of the lysine and the carboxylate residues using the hydrogen link atom scheme. The QM part was described using the B3LYP-D3(BJ)/cc-pVDZ level of theory ^45–48^. The MM region consisted of the rest of the protein and the crystallographic waters. Those were treated using the AMBER FF14SB force field ^49^ and the TIP3P model ^50^, respectively. Residues with at least one sidechain atom within 5 Å of the retinal chromophore were included in the active region and allowed to relax. The remaining protein atoms were kept fixed at their crystallographic positions.

The QM/MM optimized geometries were further utilized to compute the excitation energies using the algebraic diagrammatic construction scheme to second order with the resolution-of-identity (RI) approximation (RI-ADC(2)) ^51^, in combination with the cc-pVDZ basis set. The Turbomole software ^52^ was used for the energy computations. The vibronic lineshape was simulated using the second-order cumulant expansion method ^53^. In this approach, vibronic coupling is described via the spectral density ^54^, which is used to compute the homogeneous lineshape. To obtain the required input, ground-state molecular dynamics simulations were performed for 10 ps with time steps of 1 fs using the GFN2-xTB method as implemented in ORCA ^55^. The same QM/MM partition and active region as mentioned above was used for the dynamics. Snapshots were then extracted at 1 fs intervals, yielding a total of 10,000 configurations. Excitation energies for these snapshots were computed at the TDA-CAM-B3LYP/def2-SVP level ^56,57^ using TeraChem ^58,59^, and subsequently used to construct the vibronic spectrum. As TDDFT systematically predicted blue-shifted excitation energies relative to the ADC(2) method, the resulting spectrum was uniformly shifted by approximately 6500 cm⁻¹ to align with the experimental absorption maximum. Additional excited state (S1) relaxed potential energy scans were performed to assess the torsional flexibility of the retinal chromophore. The torsional barriers associated with rotation about the C12–C13=C14–C15 and C6–C7=C8–C9 dihedrals in NeoR were evaluated by incrementally rotating the corresponding dihedral angles in 10° steps. These scans employed the same extended QM region used for ground-state geometry optimization and excitation-energy calculations.

To rationalize the absence of 13-*cis* isomerization in NeoR, the C12–C13=C14–C15 torsional barrier was also computed for bacteriorhodopsin (PDB ID: 7Z0A ^60^). In this case, the QM region comprised the protonated Schiff base, residues D85 and D212 (both deprotonated), and the hydrogen-bonded water molecule. The Schiff base and amino acid residues were truncated between the Cα and Cβ atoms to define the QM/MM boundary.

Following the relaxed scan of the C13=C14 double bond torsion, the protein environment was removed and single point energies of the retinal chromophore were computed to model the torsional barriers without the effect of the protein. In this case, the conformation of the chromophore as obtained from the relaxed scan in the protein environment was kept intact without further optimization. Mulliken charge analysis was performed to monitor S1 charge distribution across the retinal at each step of the relaxed scan, both in the presence and absence of the protein environment. The charge analysis and single point excitation energy calculations were also computed by protonating the counterions or mutating them to alanine for both NeoR and BR, and further for mutating E141 to alanine in NeoR to explore the role of the counterion electrostatics on isomerization.

All excited-state relaxed scan and charge analysis computations were performed at (TDA)-CAM-B3LYP-D3(BJ)/cc-pVDZ level of theory using ChemShell interfaced with ORCA.

## Data availability

Data supporting the findings of this manuscript are available from the corresponding authors upon reasonable request. Atomic models from X-ray crystallography have been deposited in the Protein Data Bank (PDB) under accession codes 30BM (NeoR-NIR; 100K; rotating-crystal), 30BN (NeoR-UV, serial); 30BO (NeoR-NIR; RT; serial), 30BP (NeoR-UV; Zn^2+^; serial).

## Acknowledgements

We are deeply thankful to Dr. Gleb Bourenkov for his help in obtaining X-ray fluorescence data on *Rg*NeoR crystals. K.K. has been supported by EMBL Interdisciplinary Postdoctoral Fellowship (EIPOD4) under Marie Sklodowska-Curie Actions Cofund (grant agreement no. 847543). The work of K.K. was also supported by the European Molecular Biology Laboratory. The work was supported by the German Research Foundation (DFG) through grant 509731234 (BR 6764/2 to M.B. and BA 2242/4 to F.B.), SFB1078 “Protonation Dynamics in Protein Function”, project number 221545957, subproject B6 (to P.Hi.), and the DFG under Germany’s Excellence Strategy—EXC 311 2008/1 (UniSysCat)-390540038, Research Unit E (to P.Hi and P.He.), as well as EXC-2033-390677874 RESOLV (to I.S.). I.S. and P. He. acknowledge support by the German-Israeli-Foundation (GIF NEXUS, I-1560-207.9/2023). We further notice support by the Volkswagen Foundation (Pioneering Research, grant number 0071440-00 to I.S.), the Human Frontier Science Project (HFSP -RGP019/2024 SCHAPIRO to I.S.) and the European Research Council (ERC, Synergy “SOL” 951644, P.He.). P.He. is a Hertie Professor and is supported by the Hertie Foundation.

## Author contributions

K.K. crystallized NeoR and determined the structures of NeoR-NIR and NeoR-UV. B.B. and M.B. expressed and purified all protein samples. M.G. and F.B. acquired and analyzed the FTIR data. A.K., S.K., and P.Hi. performed and analyzed the Raman measurements. P.N. carried out and analyzed the QM/MM simulations. P.N. and I.S. interpreted the computational results and described them in the manuscript. The project was conceived and primarily supervised by M.B., together with P.He. The manuscript was written by M.B., I.S., P.He., and P.Hi., with contributions from all authors.

