## Supplementary Materials for "Structural Basis for Near Infrared Absorption and Noncanonical Photoisomerization in Neorhodopsin"

Supplementary Tables S1 to S2

Supplementary Figures S1 to S12

Supplementary References

**Table S1. X-ray crystallography data collection and structure refinement statistics.**

\*Values in parentheses are for the highest-resolution shell.

|  | NIR state | NIR state, RT | UV state | UV state, Zn bound |
| --- | --- | --- | --- | --- |
| PDB ID | 30BM | 30BO | 30BN | 30BP |
| Temperature (K) | 100 | 293 | 100 | 100 |
| Serial (yes/no) | no | yes | yes | yes |
| Crystals merged | - | 73,221 | 19,027 | 53,539 |
| Data collection |  |  |  |  |
| Space group | P2<sub>1</sub>2<sub>1</sub>2<sub>1</sub> |  |  |  |
| Cell dimensions |  |  |  |  |
| *a*, *b*, *c* (Å) | 75.8, 97.5, 109.1 | 77.7, 96.3, 113.1 | 75.8, 96.8, 108.4 | 76.0, 95.6, 108.0 |
| $\alpha$, $\beta$, $\gamma$ (°) | 90, 90, 90 | 90, 90, 90 | 90, 90, 90 | 90, 90, 90 |
| Resolution (Å) | 72.721-2.112 (2.290-2.112)* | 113.6-2.4 (2.48-2.40)* | 108.70-2.50 (2.59-2.50)* | 107.53-2.30 (2.38-2.30)* |
| R<sub>pim</sub>/R<sub>split</sub>, % | 9.6 (140.7) | 8.4 (73.4) | 12.6 (116.6) | 6.3 (81.9) |
| I/σ(I) | 7.5 (0.6) | 6.2 (0.9) | 5.7 (0.8) | 8.8 (1.2) |
| Completeness (%) | 93.0 (70.8) | 100.0 (100.0) | 100.0 (100.0) | 100.0 (100.0) |
| Redundancy | 13.6 (14.2) | 822.9 (558.0) | 380.7 (248.3) | 1036.5 (713.0) |
| CC*, % | - | 100.0 (35.0) | 99.9 (60.2) | 99.9 (43.0) |
| CC<sub>1/2</sub>, % | 99.7 (18.8) | - | - | - |
| Refinement |  |  |  |  |
| Resolution (Å) | 20-2.11 | 20-2.40 | 20-2.80 | 20-2.40 |
| No. reflections | 31,553 | 32,176 | 18,344 | 30,063 |
| R<sub>work</sub>/R<sub>free</sub> (%) | 22.8/25.9 | 22.0/23.9 | 23.9/28.2 | 24.4/25.6 |
| No. atoms |  |  |  |  |
| Protein | 4,311 | 4,263 | 4,122 | 4,136 |
| Retinal | 40 | 40 | 40 | 40 |
| Lipid | 144 | 144 | 140 | 80 |
| Solvent | 83 | 39 | 65 | 85 |
| *B*-factors (Å<sup>2</sup>) |  |  |  |  |
| Protein | 43 | 74 | 71 | 69 |
| Retinal | 40 | 62 | 72 | 63 |
| Lipid | 54 | 96 | 80 | 70 |
| Solvent | 42 | 72 | 62 | 73 |
| R.m.s. deviations |  |  |  |  |
| Bond lengths (Å) | 0.005 | 0.007 | 0.005 | 0.004 |
| Bond angles (°) | 1.552 | 1.726 | 1.416 | 0.679 |

**Table S2. QM/MM excitation energies for different protonation models of the EDE triad in NeoR computed at the RI-ADC(2)/cc-pVDZ level of theory. E141 is protonated in all cases.**

| S. No. | Charge of EDE triad | Protonation | | | $\lambda^{\max}$ | |
| --- | --- | --- | --- | --- | --- | --- |
|  |  | E136 | D140 | E262 | nm | eV |
| 1 | -3 | - | - | - | 348 | 3.56 |
| 2 | -2 | ✓ | - | - | 515 | 2.41 |
| 3 | -2 | - | ✓ | - | 549 | 2.26 |
| 4 | -2 | - | - | ✓ | 526 | 2.36 |
| 5 | -1 | ✓<sup>a</sup> | ✓ | - | 663 | 1.87 |
| 6 | -1 | ✓<sup>b</sup> | - | ✓ | 660 | 1.88 |
| 7 | -1 | - | ✓ | ✓ | 664 | 1.87 |
| 8 | 0 | ✓ | ✓ | ✓ | 641 | 1.93 |

<sup>a</sup>Proton moves to E262 during QM/MM optimization; <sup>b</sup>Proton moves to D140 during QM/MM optimization

### Supplementary Figures

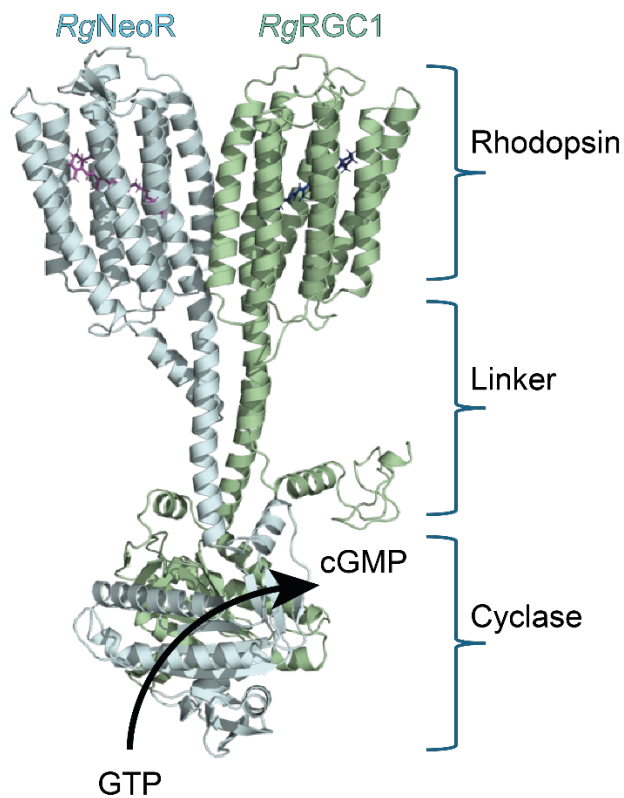

**Fig. S1. AF2-model <sup>1</sup> of a heterodimeric Rhodopsin-guanylyl cyclase from *R. globosum*.**

The two subunits, NeoR (cyan) and RGC1 (green), are depicted as cartoons, with their functional domains indicated.

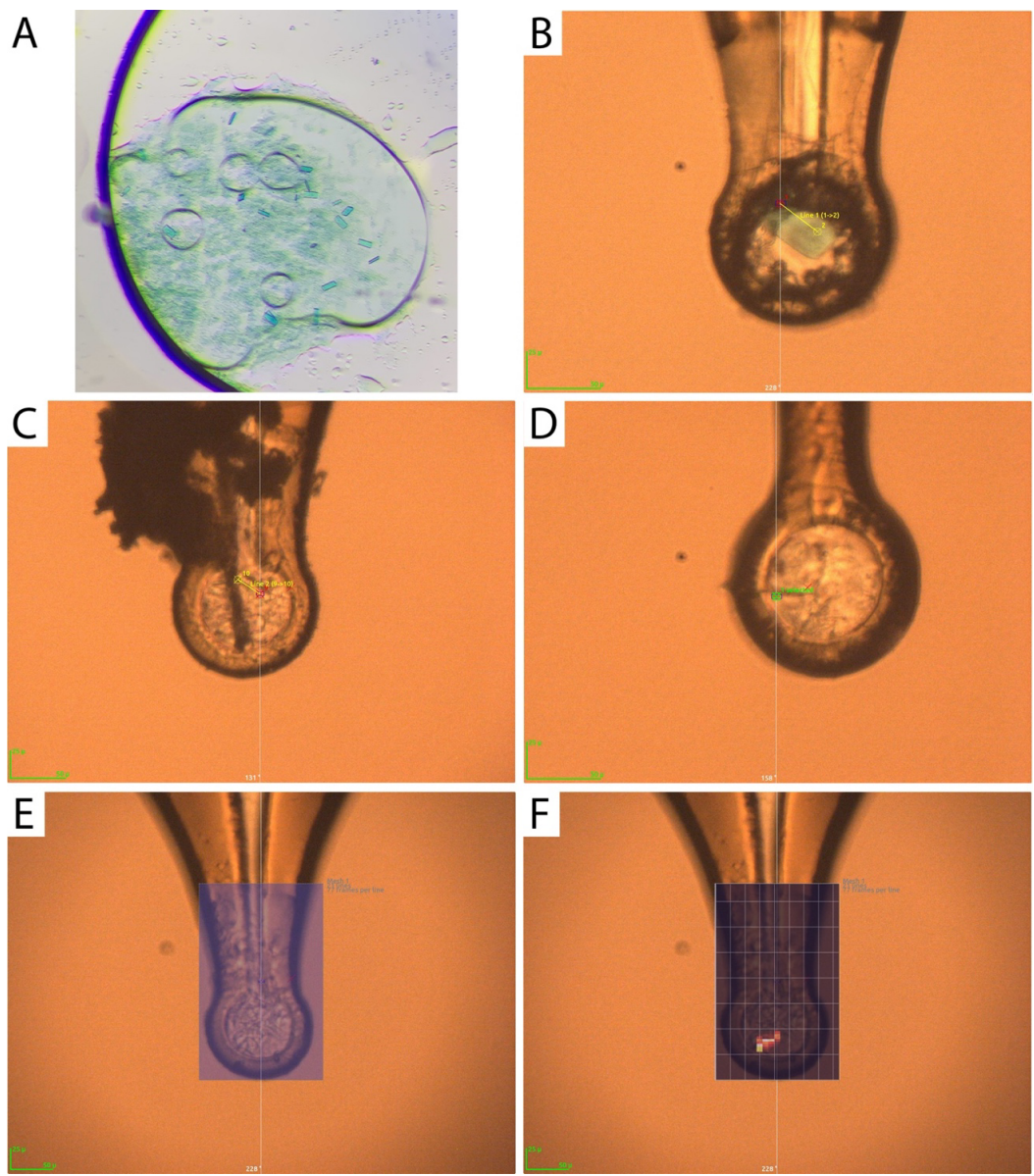

**Fig. S2. Crystals of NeoR in the NIR and UV absorbing states.**

**A.** Crystals of NeoR in the NIR-absorbing state in the crystallization probe. **B.** Snapshot of a crystal of NeoR in the NIR-absorbing state harvested in the loop and frozen at 100K mounted on the goniometer of P14 endstation of PETRAIII synchrotron. **C-D.** Snapshots of crystals of NeoR in the UV-absorbing state harvested in the loop and frozen at 100K mounted on the goniometer of P14 endstation of PETRAIII synchrotron. **E-F.** Presentation of the crystal centering procedure for data collection on NeoR in the UV-absorbing state. Crystals could not be detected from visual inspection; therefore, raster scan with microfocus X-ray beam ($5 \times 7 \mu\text{m}^2$) has been performed (panel E, blue area). Diffraction analysis indicated position of the crystal in the loop (resulting Dozor score heat map overlayed with the original view of the loop in panel F).

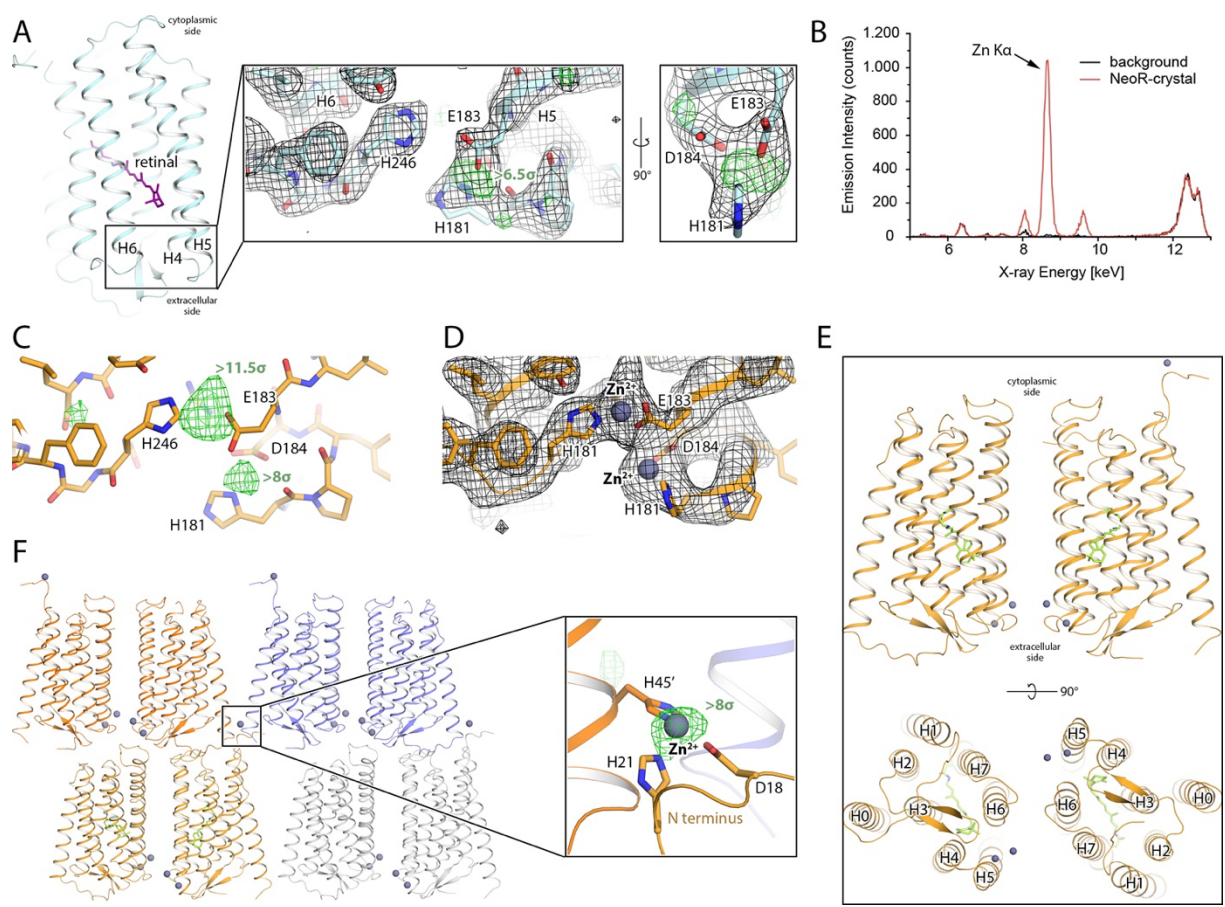

**Fig. S3. $\text{Zn}^{2+}$ binding in NeoR.**

**A.** Evidence of the ion binding in the structure of the NIR-absorbing state of NeoR. $2F_o - F_c$ electron density map is shown with black mesh and contoured at the level of $1\sigma$. $F_o - F_c$ difference electron density map is shown with black mesh and contoured at the level of $3\sigma$. **B.** X-ray fluorescence spectrum of the NeoR crystal. The main band indicates presence of $\text{Zn}^{2+}$. **C.** Examples of $F_{o\text{Zn}} - F_{o\text{noZn}}$ isomorphous difference electron density map contoured at the level of $4\sigma$ in the region between the extracellular tips of the TM helices 5 and 6. **D.** Examples of $2F_{o\text{Zn}} - F_{c\text{Zn}}$ electron density map contoured at the level of $1\sigma$ in the region between the extracellular tips of the TM helices 5 and 6. **E.** Overall side view and view from the extracellular side at the identified $\text{Zn}^{2+}$ binding sites in the NeoR dimer. **F.** $\text{Zn}^{2+}$ identified at the interlayer crystal contacts coordinated by the N terminal residues H21 and D18 and H45' of the NeoR protomer from the adjacent layer within the type I crystal.

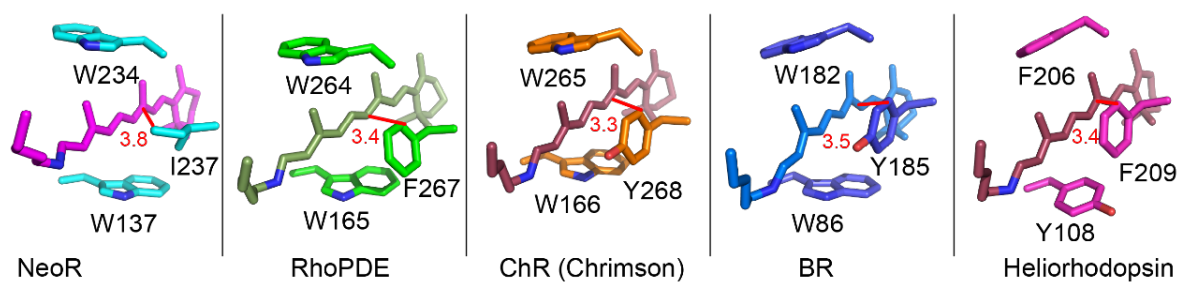

**Fig. S4. Conserved aromatic triad surrounding the retinal chromophore in various microbial rhodopsins**

(NeoR; RhoPDE, PDB: 7CJ3<sup>2</sup>; ChR, PDB: 5ZIM<sup>3</sup>; BR, PDB: 1QHJ<sup>4</sup>; Heliorhodopsin, PDB: 6SU3<sup>5</sup>). Closed distance to the retinylidene is shown in red.

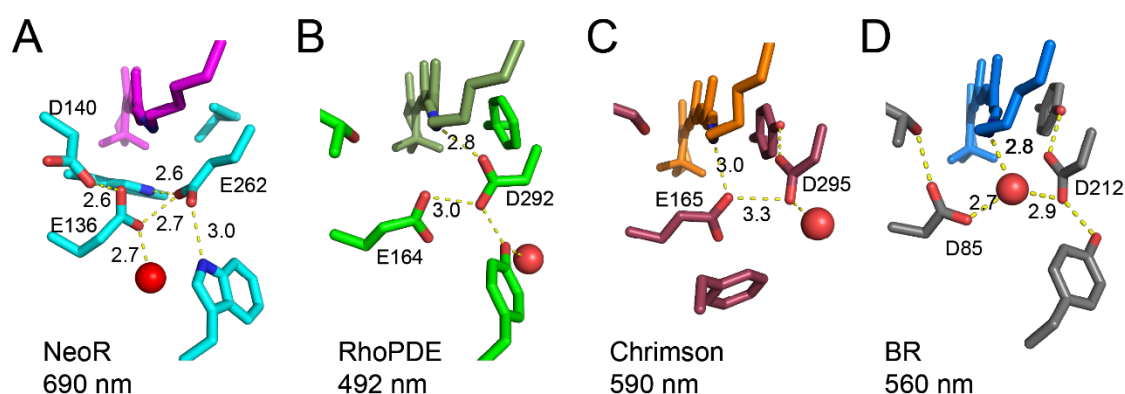

**Fig. S5. Structure of the RSB region in various microbial rhodopsins.**

**A.** NeoRhodopsin (NeoR-NIR); **B.** Rhodopsin-Phosphodiesterase (RhoPDE; PDB: 7CJ3<sup>2</sup>); **C.** Channelrhodopsin from *Chlamydomonas noctigama* (Chrimson-ChR; PDB: 5ZIM<sup>3</sup>); **D.** Bacteriorhodopsin (BR; PDB: 1QHJ<sup>4</sup>). Distances given in Å.

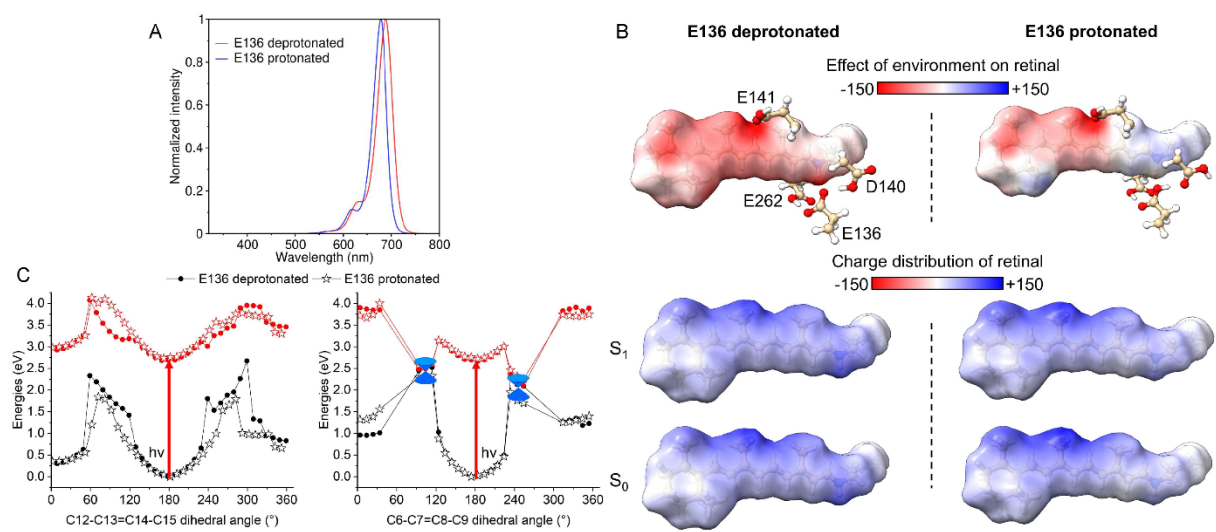

**Fig. S6. Comparison of NeoR-NIR derived QM/MM simulations with protonated or deprotonated E136.**

A) Lineshape of the visible absorption spectrum generated at the TDA-CAM-B3LYP. The calculated spectra are shifted by $\sim 6500 \text{ cm}^{-1}$ to align with the experimental maximum. The spectra are considered equal within the accuracy of the quantum chemical method. B) Upper panel: Electrostatic potential (in kT/e) of the protein projected on the surface of the retinal protonated Schiff base of NeoR-NIR. The EDE counterion triad and E141 residues are shown. Lower panel: Charges of the chromophore in its ground ($S_0$) and excited ($S_1$) states mapped on its surface. C) Relaxed potential energy scans along the C13=C14 (left panel) and C7=C8 (right panel) double-bond torsional coordinates.

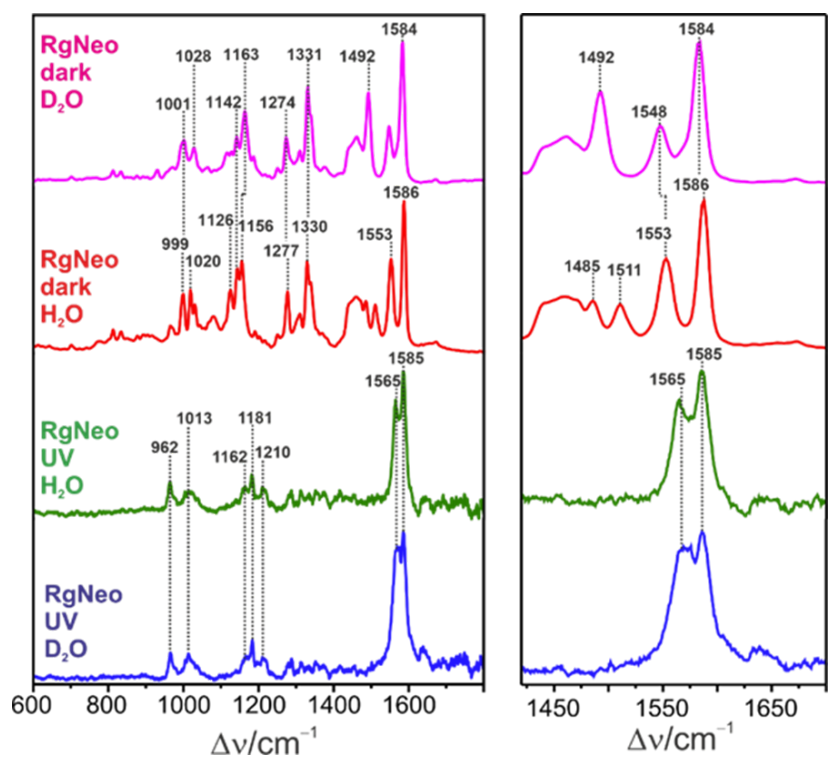

**Fig. S7. Resonance Raman spectra of the dark state NeoR-NIR in H<sub>2</sub>O (red) and D<sub>2</sub>O (magenta).**

Spectra are compared with the photoinduced NeoR-UV state in H<sub>2</sub>O (green) and D<sub>2</sub>O (blue). All spectra were measured at 90 K. The spectra of the dark state, obtained with 1064 nm excitation, show a large number of spectral changes due to the replacement of the RSB proton by a deuteron. In contrast, the spectra of the NeoR-UV are, within the S/N ratio, the same in H<sub>2</sub>O and D<sub>2</sub>O, due to the lack of an exchangeable proton of the chromophore. Thus, in contrast to the NeoR-NIR state, the UV-state includes a deprotonated RSB. The 1585 $\text{cm}^{-1}$ is tentatively attributed to the C=C stretching of the retinal lacking the Schiff base linkage to the protein <sup>6</sup>. The hydrolysis of the Schiff base bond in the protein is reminiscent of the transient retinal formation in the long-lived intermediate P353 of the Channelrhodopsin variant ChR-C128T <sup>7</sup>.

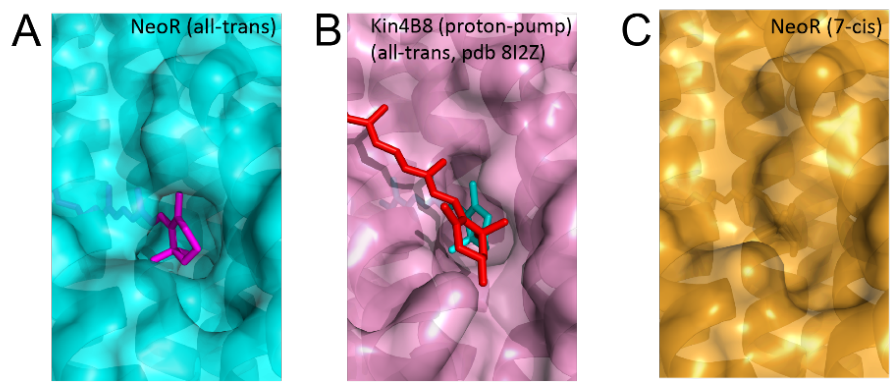

**Fig. S8. Fenestration towards the retinal binding pocket.**

NeoR-NIR (all-*trans* form). B) Xanthorhodopsins Kin4B8 (all-*trans*) with bound zeaxanthin (red; PDB: 8I2Z<sup>8</sup>). C) NeoR-UV (7-*cis* form).

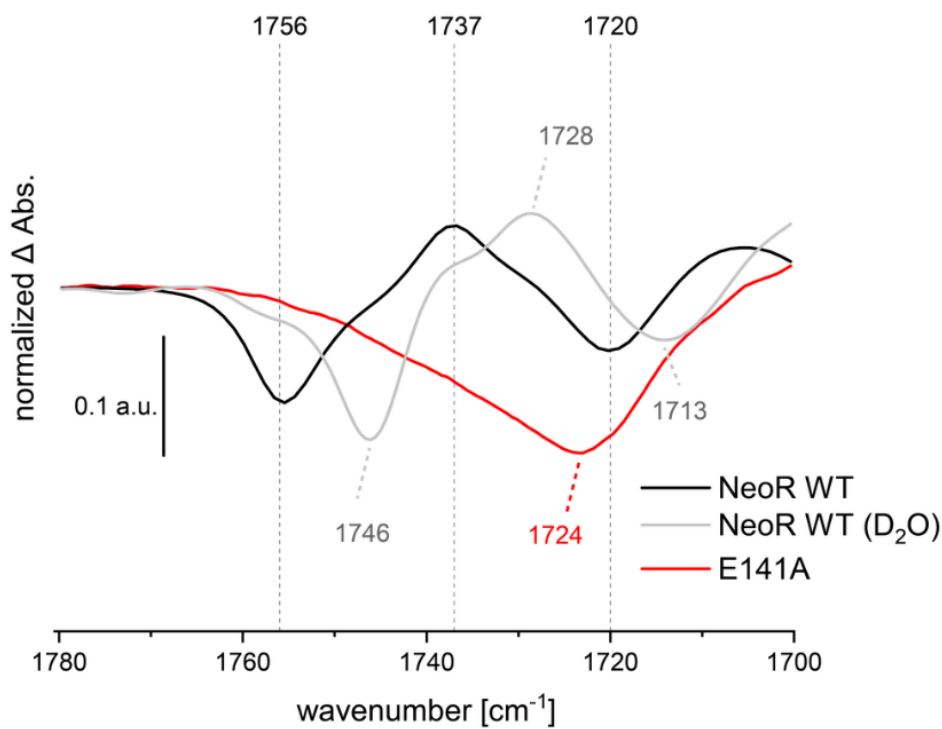

**Fig. S9. NeoR-UV minus NeoR-NIR FTIR difference spectrum in the spectral region between 1780 and 1700 $\text{cm}^{-1}$.**

**Black line:** Light-induced FTIR difference spectrum (UV-minus-NIR state) of the NeoR wild type (WT) used for crystallography, closely resembling that reported for the ortholog OmNeoR<sup>9</sup>. **Grey line:** Light-induced FTIR difference spectrum (UV-minus-NIR state) of the NeoR wild type in $\text{D}_2\text{O}$. **Red line:** Light-induced FTIR difference spectrum (UV-minus-NIR state) of the NeoR E141A mutant. The band pattern at $1756\text{ cm}^{-1}$ (–) / $1737\text{ cm}^{-1}$ (+) in the wild type, which is downshifted to $1746\text{ cm}^{-1}$ (–) / $1728\text{ cm}^{-1}$ (+) in $\text{D}_2\text{O}$, indicates a light-induced change in the hydrogen-bonding environment of protonated E141 during the transition from the NeoR NIR dark state to the NeoR UV state. This band pattern is absent in the E141A mutant, allowing an unambiguous assignment to E141. The residual negative band in the E141A difference spectrum is tentatively assigned to D140.

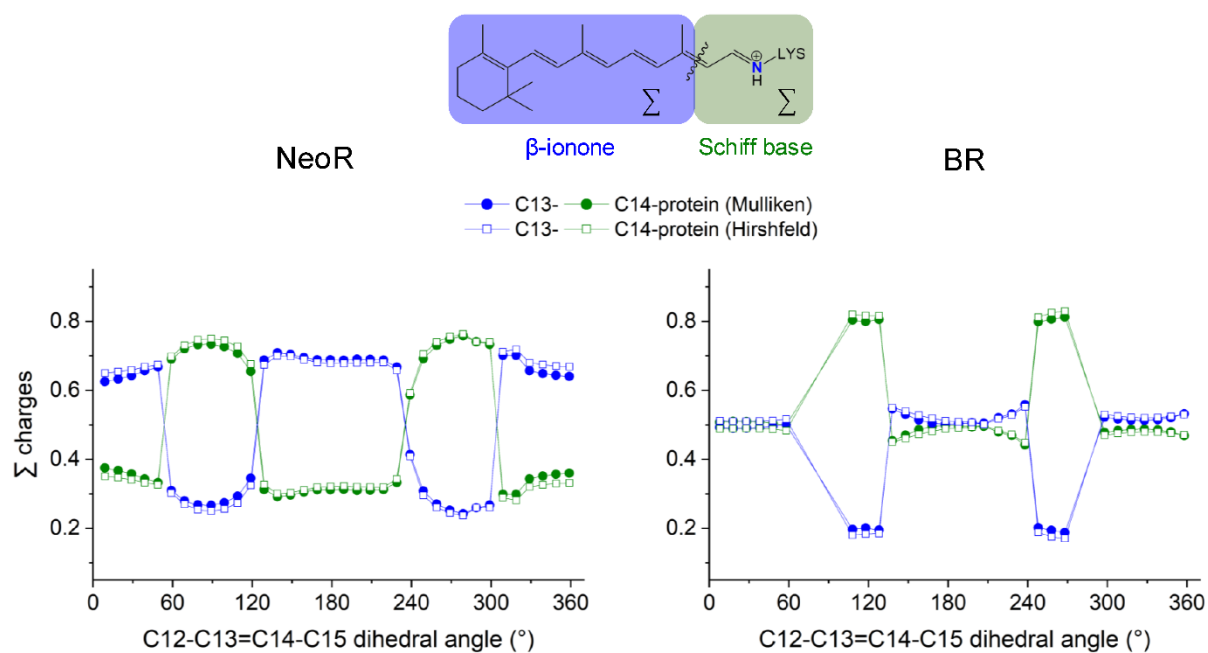

**Fig. S10. Comparison of Mulliken or Hirshfeld based charge distribution.**

Excited (S1) state charge distribution analysis based on Mulliken and Hirshfeld charges along the Schiff base and β-ionone fragments of the retinal chromophore during the relaxed scan in NeoR and BR.

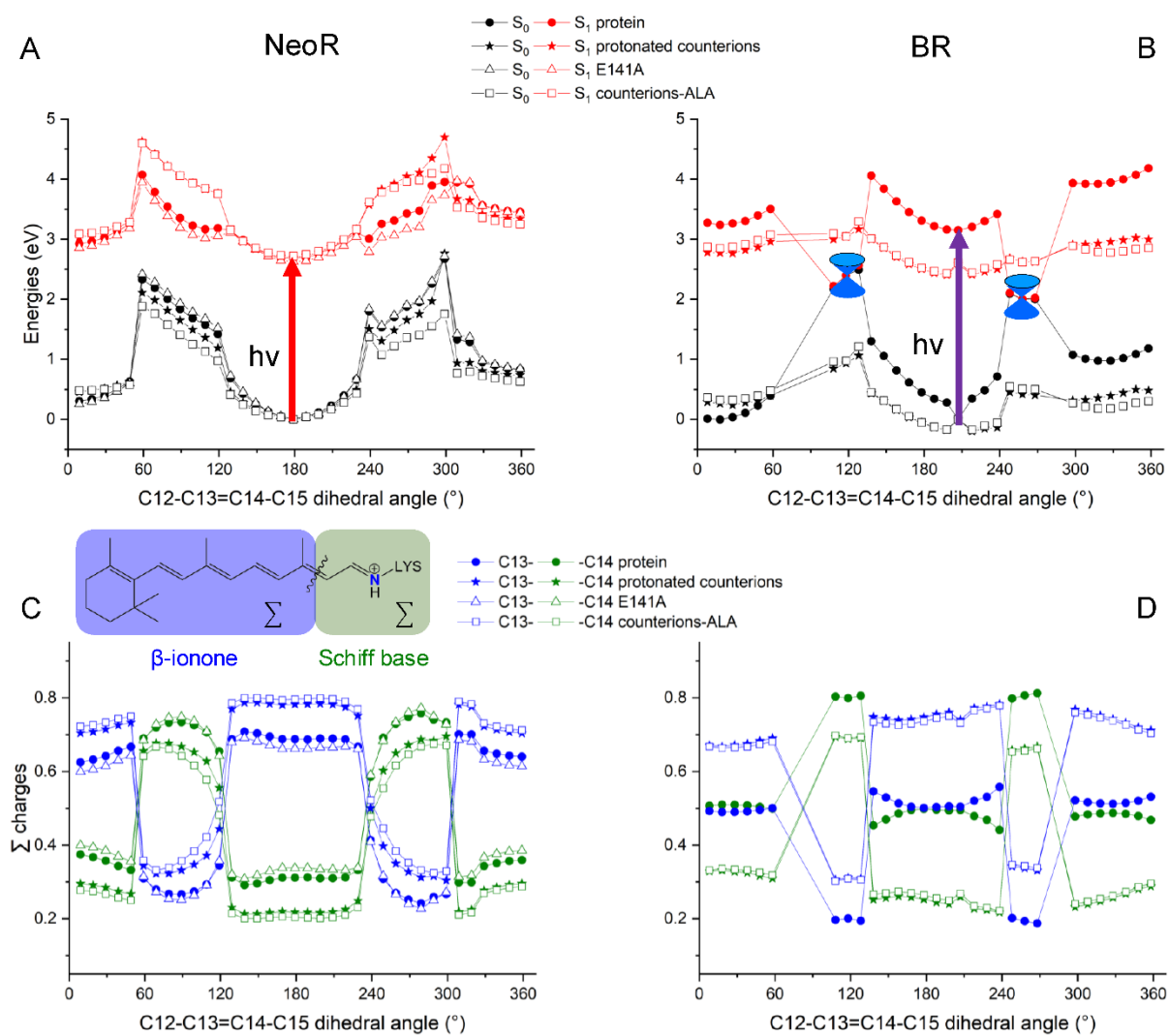

**Fig. S11. Relaxed potential energy scans of NeoR-NIR and BR.**

(A, B) Relaxed potential energy scans along the C13=C14 double-bond torsional coordinate were performed for models containing either one (NeoR-NIR) or two (BR) negative counterions. Additional simulations were conducted with fully protonated counterions and mutating the counterions to ALA for both NeoR and BR, as well as for the NeoR E141A mutant. (C, D) Excited ($S_1$) state charge distribution (Mulliken analysis) along the Schiff base and β-ionone fragments of the retinal chromophore during the relaxed scan in the two proteins, respectively.

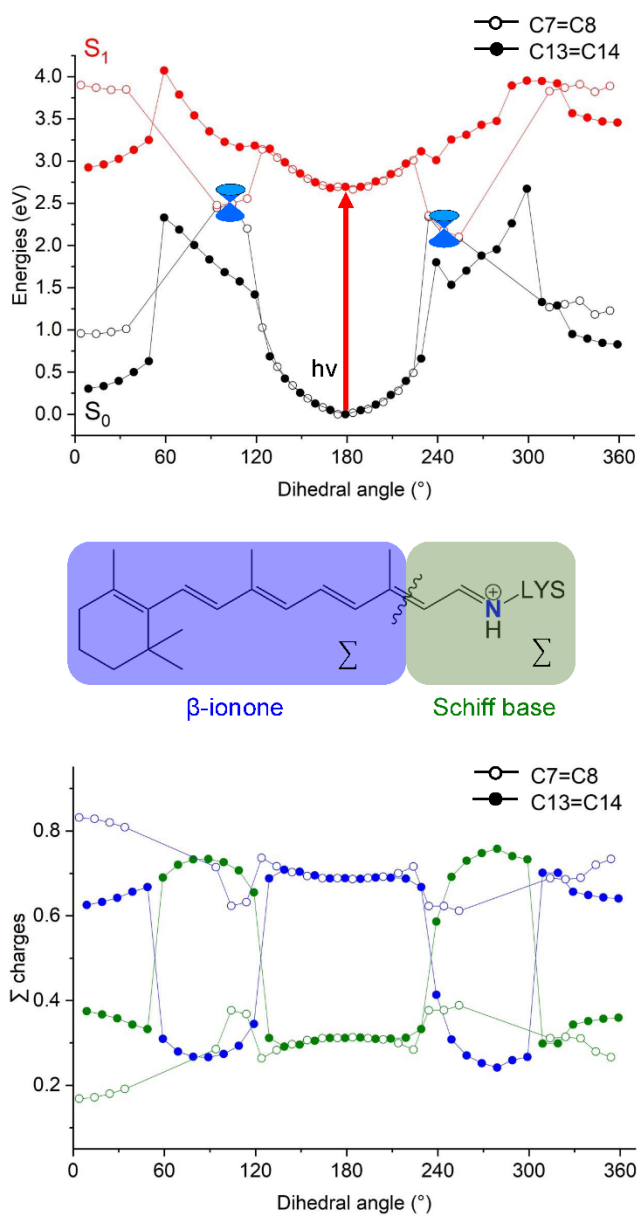

**Fig. S12. Relaxed potential energy scans and charge distribution of NeoR-NIR.**

Upper panel: Relaxed potential energy scan along the C7=C8 and C13=C14 double-bond torsional coordinates for NeoR with one negative counterion (E136). Lower panel: Excited (S<sub>1</sub>) state charge distribution (Mulliken analysis) along the Schiff base and β-ionone fragments of the retinal chromophore during the relaxed scan.
